# CBFβ initiates the hematopoietic stem cell program without obligatory binding to RUNX

**DOI:** 10.1101/172957

**Authors:** Aldo Ciau-Uitz, Philip Pinheiro, Arif Kirmizitas, Claire Fernandez, Roger Patient

## Abstract

Hematopoietic stem cells (HSCs) emerge from hemogenic endothelium (HE) localised in the embryonic dorsal aorta (DA). Here we show that Runx1, a transcription factor essential for HSC emergence, controls HE establishment in the absence of its non-DNA-binding partner, CBFβ, and that a CBFβ-binding-deficient Runx1 mutant form can activate the HE program in the DA. Nevertheless, CBFβ is also essential for HSC emergence by regulating the specification of definitive hemangioblasts (DHs), the precursors of the DA and HE, in the lateral plate mesoderm where it mediates VEGFA induction by BMP signalling. Surprisingly, no Runx gene is expressed in DHs and the pharmacological inhibition of CBFβ binding to Runx is not detrimental for DH, confirming that CBFβ functions independently of Runx. Thus, we have uncovered, for the first time, that CBFβ regulates gene expression without Runx, breaking the dogma in which CBFβ ‘s gene regulatory functions are strictly dependent on its binding to Runx.

**HIGHLIGHTS:**

- Runx1 and CBFβ play independent roles in the establishment of the HSC lineage
- Runx1 binding to CBFβ is not required for HE establishment
- CBFβ is downstream of BMP and regulates endogenous VEGFA expression in DH
- Binding to Runx is not obligatory for CBFβ function

## INTRODUCTION

Core binding factor (CBF), is a heterodimeric transcription factor (TF) complex composed of one of the three runt-related DNA-binding α subunits (Runx1, Runx2, Runx3) and the non-DNA-binding β subunit (CBFβ). The α subunits are capable of binding to DNA alone albeit at lower affinities compared to when complexing with CBFβ (Tang et al., 2000). Additionally, CBFβ confers protein stability to the α subunits by preventing their ubiquitination and degradation by the proteasome (Huang et al., 2001; Park et al., 2016; Qin et al., 2015). To date, CBFβ’s transcriptional activity appears to be completely dependent on its physical interaction with the α subunits, i.e. its interaction with Runx is obligatory.

Runx proteins act as master regulators in different cell lineages with some overlapping functions. Of these, Runx1 has been established as a key regulator of hematopoiesis; in particular, Runx1 is essential for HSC generation. Runx1 is also frequently altered in chromosomal translocations associated with human leukemia (Chin et al., 2015) and is a critical regulator of haematopoietic-specific genes in a variety of blood cells. Runx1 expression during embryogenesis is prominent in the developing haematopoietic system and is needed in several hematopoietic processes such as megakaryocytic differentiation, erythropoiesis, B- and T-cell development; and, critically, the establishment of definitive hematopoiesis. HSCs are at the foundation of definitive hematopoiesis and they are transiently generated in the aorta-gonads-mesonephros (AGM) region of the midgestation embryo. Recent live imaging studies have demonstrated that HSCs are directly generated from a small cohort of HE cells localised in the ventral wall of the DA through a process termed as an endothelial to hematopoietic transition (EHT)(Bertrand et al., 2010; Boisset et al., 2010; Kissa and Herbomel, 2010). Through conditional deletion, it has been demonstrated that Runx1 is specifically required for the generation of HSCs but not thereafter, indicating that it is an essential regulator of EHT (Chen et al., 2009). Indeed, EHT in the zebrafish DA fails in the absence of Runx1 (Kissa and Herbomel, 2010). However, how Runx1 controls EHT is yet to be elucidated.

The deletion of either Runx1 or CBFβ gives nearly identical hematopoietic phenotypes in the mouse, consistent with CBFβ function being dependent on Runx1. Importantly, CBFβ-deficient mice die at midgestation due to a lack of fetal liver hematopoiesis, a phenotype reminiscent of Runx1 deficiency, indicating that CBFβ is required for definitive hematopoiesis (Niki et al., 1997; Okuda et al., 1996; Wang et al., 1996). In agreement, using a CBFβ-GFP, it has been determined that all c-kit^hi^ cells from the fetal liver and the AGM, which contain HSCs and nascent HSCs respectively, express CBFβ (Kundu et al., 2002). Although CBFβ function in HSC emergence has been minimally characterized, some lines of evidence indicate that it is required. In embryos expressing a knocked-in CBFβ-MYH11, a fusion protein which acts in a dominant negative manner and therefore abolishes CBFβ function, the population of c-kit^hi^ cells is absent in the AGM (Kundu et al., 2002), indicating that HSC generation is impaired. The Ly6a regulatory sequences drive expression in HE and all functional HSCs in the DA (de Bruijn et al., 2002) and the generation of HSCs in CBFβ-deficient embryos can be rescued by the expression of a GFP/CBFβ fusion protein driven by the Ly6a regulatory sequences, confirming that CBFβ is essential for EHT (Chen et al., 2011). Controversially, the emergence of hematopoietic stem/progenitor cells (HSPCs) in zebrafish CBFβ mutants appears normal but definitive haematopoiesis fails because these cells cannot migrate to the fetal liver equivalent, the caudal hematopoietic tissue (Bresciani et al., 2014). The reasons for this discrepancy have not been investigated; nevertheless, this study provides further evidence that Runx1 can function independently of CBFβ.

In contrast to Runx1, which has a minimal impact on the expansion and maintenance of HSCs, the requirement for CBFβ appears to continue after the emergence of HSCs (Cai et al., 2011; Miller et al., 2001; Wang et al., 2015). On one hand, this may suggest the involvement of either Runx2 or Runx3. Indeed, Runx3 and Runx1 have been reported to play redundant roles in the maintenance of HSCs (Wang et al., 2015). Interestingly, Runx3-deficiency in the zebrafish causes a reduction in the number of emerging HSPCs (Kalev-Zylinska et al., 2003), suggesting that Runx3 is involved in the expansion or maintenance of nascent HSCs. On the other hand, although less likely, this may suggest that CBFβ has Runx-independent functions, however, a Runx1-independent role for CBFβ in the emergence and maintenance of HSCs remains to be identified.

The emergence of HSCs in the mouse AGM has mainly been studied between stages E10.5-11.5, when functional HSCs can be detected and intra-aortic clusters (IACs) are in the process of budding or have already budded from HE. Thus, these studies analyse the mechanism of the last step in the emergence of HSCs, EHT, rather than HE establishment. It has been suggested that establishment of HE in the mouse DA initiates as early as stage E8.5 (Swiers et al., 2013), but very little is known of the processes taking place before EHT. Here, taking advantage of the accessibility of the Xenopus embryo before EHT, we investigate the role of Runx1 and CBFβ in the establishment of HE and show that Runx1 expression in the DA initiates before the onset of circulation whereas CBFβ initiates expression later, at the beginning of IAC budding. Furthermore, we show that Runx1 establishes the HE program in the absence of CBFβ and without perturbing arterial and venous specification. We also show that HE is not established in CBFβ-deficient embryos because CBFβ is expressed in, and essential for, the specification of their lateral plate mesoderm (LPM) precursors, DHs. Furthermore, we show that CBFβ is downstream of BMP signaling and is essential for VegfA expression in DHs and does so in the absence of Runx proteins. Thus, for the first time, we have unveiled a role for CBFβ which is independent of its binding to Runx, breaking the dogma in which CBFβ regulates gene expression strictly by physically interacting with Runx.

## RESULTS

### Runx1 expression in the DA initiates before lumenisation and coincides with the loss of the venous program

Expression of Runx1 defines the hemogenic potential of endothelial cells (Chen et al., 2009). Using Runx1 expression as an indicator, we first determined when HE is established within the HSC lineage. Previously, we showed that Runx1 is not expressed in LPM mesoderm, where the precursors of the DA, DHs, are localised (Ciau-Uitz et al., 2013). From the LPM, a subpopulation of DHs migrate to the midline where they coalesce to form the DA (Cleaver and Krieg, 1998). As they migrate, DHs do not express hematopoietic genes (Ciau-Uitz et al., 2000). Runx1 expression was first detected in the DA by in situ hybridization (ISH) at stage 35 (Figure 1A), this is just after the heart initiates beating but before DA lumenisation (stage 37) and the onset of blood circulation (stage 38/39). Runx1 expression analysis on sectioned embryos shows that its expression is restricted to the ventral wall of the lumenised DA (Figure 1B). Initially, these cells show an elongated, endothelial-like morphology but, with time, they undergo a morphological change so they have become rounded before they bud out to form the first IACs at stage 42 (Figure 1B). This morphological change has also been described in the mouse DA and has been suggested to be indicative of the acquisition of hemogenic potential (Boisset et al., 2010; Bos et al., 2015; Mizuochi et al., 2012).

**Figure 1.**
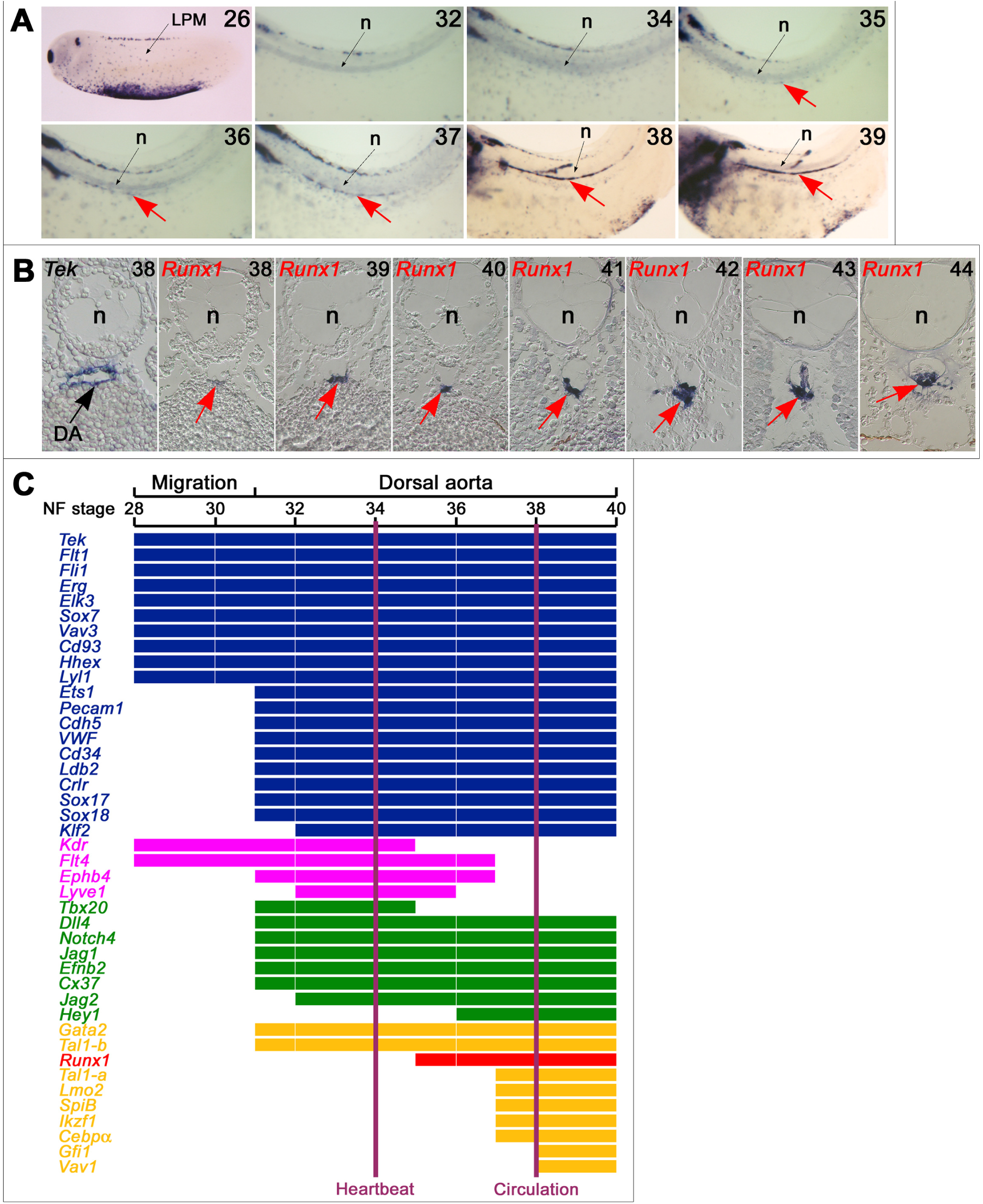
Timing of Runx1 expression and HE establishment in the DA. (**A**)Timing Runx1 expression in the DA (arrows) by ISH. Images show details of the trunk region with head to the left and dorsal to the top. The position of the LPM and notochord (n) are indicated. Numbers in top right corner indicate stage of development. (**B**)ISH on sectioned embryos showing that Runx1 expression is restricted to the ventral wall of the DA (arrows). A section stained for Tek is presented for comparison. n, notochord. Sections are transversal and 10 μ thick. Dorsal is to the top. Numbers in top right corner indicate stage of development. (**C**)Diagram summarizing the expression of hematopoietic (orange), pan endothelial (dark blue), arterial (green) and vein-affiliated (pink) genes in the DA at the time HE is established. Runx1 is shown in red. Vertical lines indicate the time when heartbeat initiates and the onset of circulation.

To better understand how HE is established in the DA, we first analysed by ISH the expression of hematopoietic genes before the first IACs sprout (stage 42) and determined that Gata2, Tal1-a, Tal1-b, Lyl1, Lmo2, SpiB, Gfi1, Ikzf1, Vav1 and Cebpα were expressed whereas Gata1, Gata3, Etv2, Etv6, PU1, SpiC, Gfi1b, aMyb, bMyb, cMyb, Mecom (Evi1), CD41, CD45, Prom1 and CBFβ were not expressed (Figure 1C and S1). With the exception of Gata2, Lyl1 and Tal1-b, all hematopoietic TFs are temporally downstream of Runx1 (Figure 1C) and potentially regulated by it. The earlier expression of Gata2 and Tal1-b is in agreement with their role as Runx1 regulators. Interestingly, only one of the Tal1 genes, Tal1-b, is upstream of Runx1. This situation is similar in the zebrafish where Scl-β is upstream while Scl-α is downstream of Runx1 and play distinct roles in HSC emergence (Zhen et al., 2013). This suggests that Tal1-a and Tal1-b may also have distinct roles in Xenopus. Regarding CBFβ, its expression in the DA coincides with the budding of the first IAC (stage 42, Figure 2A), suggesting that it plays a role in this process.

**Figure 2.**
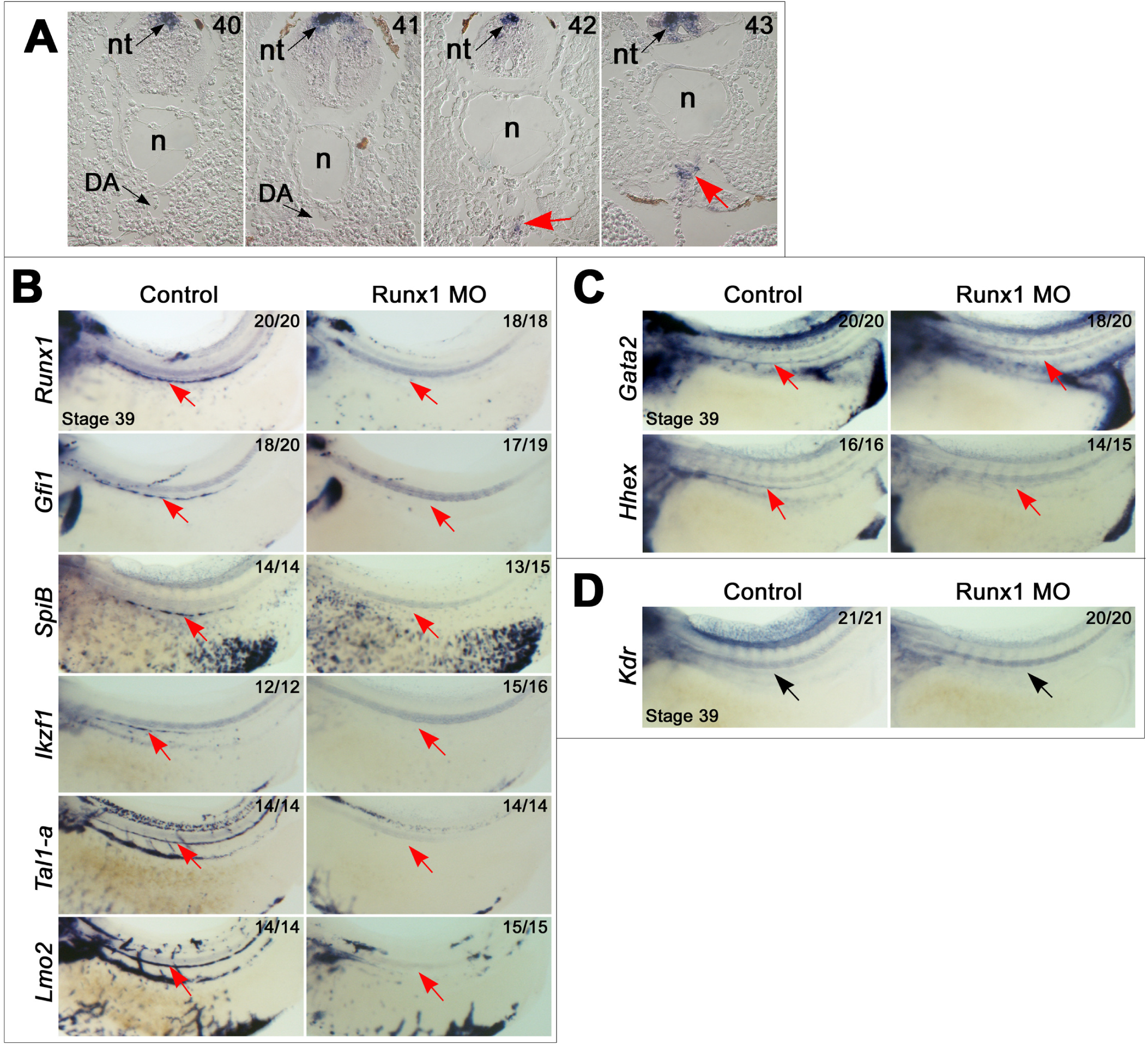
Runx1 regulates hematopoietic gene expression in the DA before the onset of CBFβ expression. (**A**)ISH on sectioned embryos showing timing of CBFβ expression in the DA (red arrows). Expression in the neural tube (nt) is indicated, n, notochord. Sections are transverse and 10μm thick. Dorsal is to the top. Numbers in top right corner indicate stage of development. (**B**)ISH at stage 39 showing that Runx1 is required for hematopoietic expression in DA. Expression of Tal1a and Lmo2 in the posterior cardinal vein (PCV) represents circulating cells and their absence in morphants suggest that the onset of circulation is delayed in this embryos. (C) ISH at stage 39 showing that the expression of blood genes expressed before the onset of Runx1 become dependent on Runx1. (**D**)ISH at stage 39 showing that the expression of vein-affiliated genes is unaffected in the DA of Runx1 morphants. Arrows in (**B**-**D**) indicate expression in the DA. Numbers in top right corner indicate proportion of embryos displaying the phenotype.

Next we analysed the expression of arterial (Notch4, Dll4, Jag1, Efnb2 and Cx37), vein (Kdr, Flt4, Lyve1 and Ephb4) and pan-endothelial genes (Tek, Flt1, Fli1, Ets1, Erg, Elk3, Hhex, CD93, CD34, Ldb2, Pecam1, VWF and Cdh5), and the endothelial expressed SoxF genes (Sox7, Sox17 and Sox18). Some pan-endothelial and vein-affiliated genes were expressed from the DH stage (Tek, Flt1, Fli1, Erg, Elk3, Hhex, Kdr and Flt4) but the genes typically regarded as expressed in mature endothelial cells (Pecam1, VWF and Cdh5) were activated after migration, upon arrival at the midline. Regarding arterial-affiliated genes, they were all activated upon arrival at the midline, reinforcing the view that the angioblasts in the DA are no longer hemangioblasts but endothelial cells. Furthermore, Etv2, an ETS TF expressed in nascent angioblasts and DHs, is not detectable by ISH in the DA (Figure S1). SoxF genes have been implicated in arterial specification and hematopoiesis, specifically in the regulation of Dll4 (Clarke et al., 2013; Corada et al., 2010); accordingly, Sox7, Sox17 and Sox18 are co-expressed with Dll4 in the DA. It is also important to note that the DA endothelium initially expresses both the arterial and vein programmes and that the vein program is lost concomitantly with the onset of Runx1 expression (Figure 1C). The markers of hematopoietic commitment, CD41 and CD45, are not expressed before stage 42 (Figure S1), indicating that, up to this stage, Runx1 expressing cells have not yet committed to the hematopoietic lineage.

### Runx1 is essential for haematopoietic expression in the DA

Although both Runx1 and CBFβ are required for the emergence of HSCs in the mouse DA, only the expression of Runx1 has been examined in detail (Swiers et al., 2013) because CBFβ is considered to be ubiquitously expressed. Our gene expression analysis shows that their expression in the DA initiates at different stages: while CBFβ expression is first detected at the stage when the first IACs are visualised (Stage 42, Figure 2A), the onset of Runx1 expression takes place much earlier, before the DA is lumenised (Figure1). Albeit with lower affinity, in vitro assays indicate that Runx1 can bind DNA in the absence of CBFβ (Tang et al., 2000), suggesting that Runx1 could regulate gene expression without CBFβ in vivo. Recent studies in the zebrafish, where the physical interaction between Runx1 and CBFβ was blocked with a small molecule, also suggest that Runx1 may regulate gene expression without CBFβ (Bresciani et al., 2014). Interestingly, known direct transcriptional targets of Runx1 such as Gfi1, SpiB (the functional equivalent of PU1 in Xenopus) and Vav1 (Denkinger et al., 2001; Huang et al., 2008; Lancrin et al., 2012; Okada et al., 1998), are expressed soon after Runx1 initiates expression in the DA (Figure 1C), suggesting that Runx1 is transcriptionally active in the absence of CBFβ. To test this, gene expression in the DA was analysed in Runx1 deficient embryos generated by the co-injection of two morpholino antisense oligonucleotides (MOs) which block the translation of the two protein forms of Runx1, P1 and P2, generated by the usage of alternative promoters (Figure S2A-C). Indeed, the expression of these genes was absent in the DA of Runx1 morphants (Figure 2B) and consequently a lack of IAC budding in the DA (Figure S2D). Additionally, the expression of Tal1-a, Lmo2 and Ikzf1 was also absent in these morphants (Figure 2B). Interestingly, expression of blood genes, whose expression initiates before the onset of Runx1 in the DA (such as Gata2, Hhex and Tal1-b), is also absent in Runx1 morphants at stage 39, indicating that their expression becomes dependent on Runx1 (Figures 2C). Furthermore, expression of Runx1 itself is missing when it is depleted, suggesting an auto regulatory loop (Figure 2B).

Runx1 has been reported to repress Kdr expression in differentiating ES cells and endothelial cell lines (Hirai et al., 2005). Interestingly, the onset of Runx1 expression coincides with the downregulation of Kdr expression in the DA (Figure 1C), suggesting that Runx1 might repress Kdr to establish the HE program. However, Kdr expression in the DA of Runx1 morphants was unaltered rather than upregulated (Figure 2D), indicating that Kdr is repressed by a Runx1 independent mechanism. Similar to Kdr, the downregulation of vein-affiliated genes (Flt4, Ephb4, Lyve1) in the DA also coincides with the onset of Runx1 expression (Figure 1C) but their expression is unaltered in Runx1 deficient embryos (Figures 2D), indicating that Runx1 is not involved in the repression of the vein programme in the DA. Furthermore, Runx1 depletion did not alter the expression of arterial and pan-endothelial genes (Figure S2E), indicating that Runx1 is specifically required for the establishment of the HE programme.

Because it has recently been argued that MOs may have non-specific effects in the zebrafish (Kok et al., 2015), the hematopoietic, arterial and vein phenotypes were confirmed by TALEN-mediated loss of function (Figures S3). Taken together, these results strongly suggest that Runx1 is transcriptionally active in the absence of CBFβ.

### Runx1 establishes HE independently of CBFβ

Having shown that CBFβ mRNA is not detected in the DA at the time Runx1 begins to establish the HE program, we sought to determine whether it is also absent at the protein level. With this aim, DA explants were dissected out from wild type embryos (Figure 3A) and subjected to western blotting using a polyclonal anti-Human CBFβ antibody. Results show that CBFβ protein is not detected in the DA at the stages of development analysed (Figure 3B) and, therefore, is not involved in the earliest stages of HE programing.

**Figure 3.**
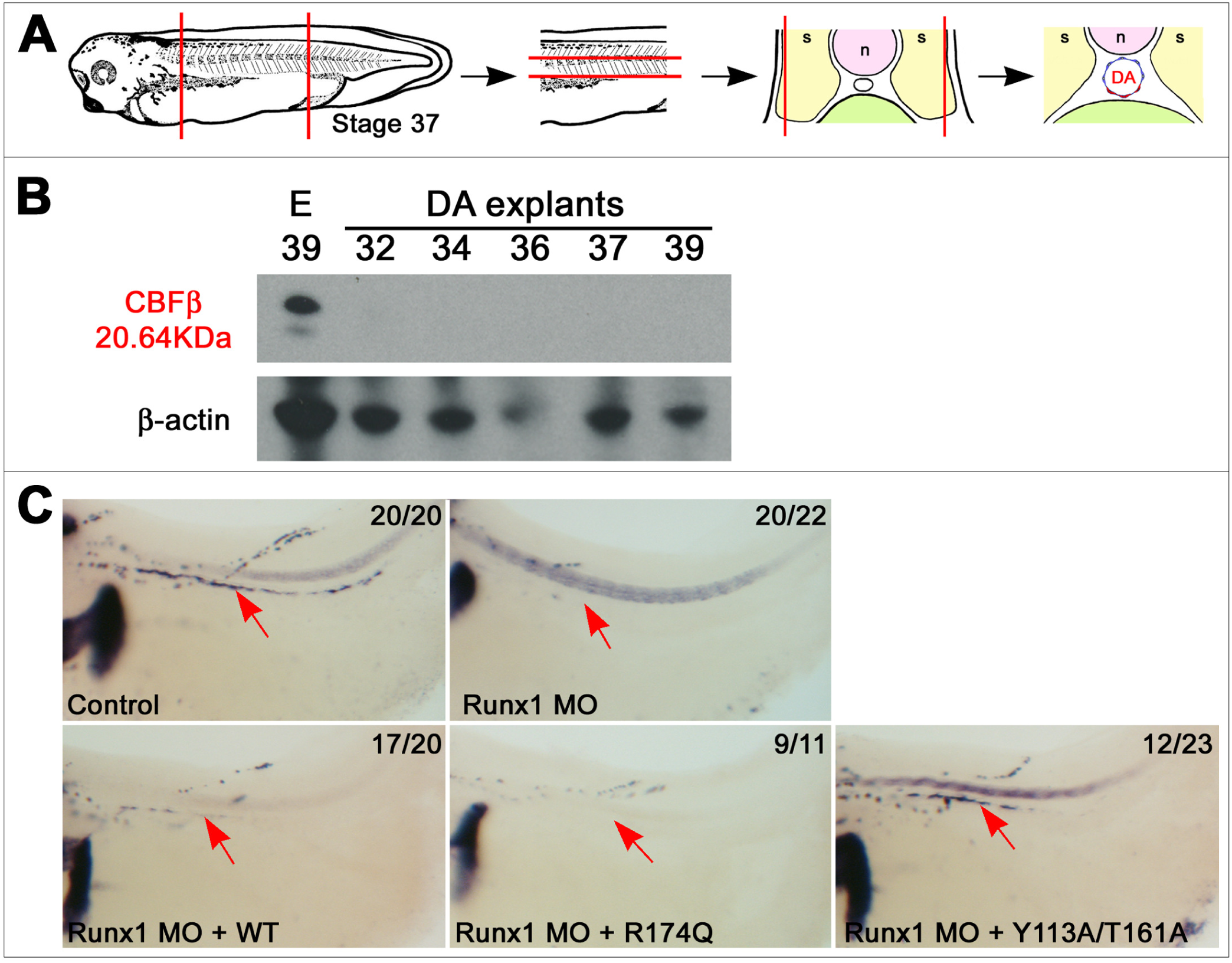
Runx1 activates gene expression in the DA in the absence of CBFβ. (**A**)Schematic of DA explant dissection for western blot analysis. (**B**)Western blot showing that CBFβ protein is not detected in the DA at the time Runx1 is transcriptionally active. Numbers indicate stage of development. E, whole embryo extract. β-actin was used as a loading control. (C) Expression of Gfi1 in the DA of Runx1 morphants is rescued with the wild type (WT) and CBFβ binding mutant (Y113A/T161A) forms of mouse Runx1 but not with a DNA-binding mutant (R174Q) form. Arrows indicate expression in the DA. Numbers in top right corner indicate proportion of embryos displaying the phenotype.

Next, we investigated whether Runx1 is sufficient for the initiation of the HE program in the DA by rescuing the Runx1 morphants with mouse Runx1 (mRunx1). Because exogenous mRNA injected into Xenopus embryos is only stable for up to 2 days (Walmsley et al., 2002) and HE starts to be established by day 4 of development, the Runx1 phenotype was rescued by transient transgenesis mediated by tol2 transposition (Urasaki et al., 2006). Thus, embryos were injected with MOs together with the tol2 vectors for transgenesis and tol2 transposase mRNA. Injection of MOs alone abolished the expression of Gfi1 in 20 out of 22 embryos (Figure 3C). As expected, Gfi1 expression was rescued in 17 out of 20 embryos coinjected with wild type mRunx1 (Figure 3C). Structural and mutagenesis analyses have revealed the amino acid residues in Runx1 which are responsible for its binding to DNA as well as those required for its dimerization with CBFβ. Thus, it has been demonstrated that an R174Q substitution in Runx1 causes a greater than 40,000-fold reduction in DNA binding without perturbing its heterodimerization with CBFβ, its structure or its thermodynamic stability while the double mutation Y113A/T161A causes a 430-fold reduction in CBFβ binding without having an effect on DNA binding (Matheny et al., 2007; Roudaia et al., 2009). The R174Q mRunx1 mutant was unable to rescue the expression of Gfi1 in Runx1 deficient embryos (Figure 3C), indicating that Runx1 DNA binding is required for the establishment of the HE program in the DA. In contrast, the Y113A/T161A double mutant rescued Gfi1 expression in 50% of the embryos analysed (Figure 3C), indicating that CBFβ is not required for Runx1 transcriptional activity in the DA. Taken together, we conclude that, in vivo, Runx1 is transcriptionally active in the absence of CBFβ and that it is necessary and sufficient for the establishment of the HE programme in the DA.

### CBFβ is required for HE specification

To confirm that CBFβ is not required for the initiation of the HE program in the DA, MOs blocking its translation were designed (Figure 4A-B) and injected into embryos. Surprisingly, the expression of Runx1, its targets (Gfi1, SpiB) and other hematopoietic genes (Gata2, Tal1-a, Lmo2), were absent in the DA of CBFβ morphants (Figure 4C), indicating that CBFβ is required for HE specification. Furthermore, the expression of Gata2 and Tal1-b, which normally initiate before the onset of Runx1 expression in the DA, was also abrogated (Figure 4D), suggesting that CBFβ acts upstream of Runx1 in HE programming. Additionally, Kdr expression in the DA was upregulated whereas the expression of arterial genes such as Notch4, Cx37, Efnb2 and Jag1, were downregulated (Figure 4E), suggesting that CBFβ is involved in the repression of the venous program as well as in the arterial specification of the DA. Because the expression of many of these genes initiates as the DA precursors coalesce in the midline at stage 31 (Figure 1C) and CBFβ protein is not detected in stage 32 DA explants (Figure 3B), these results indicate that CBFβ is required before the formation of the DA.

**Figure 4.**
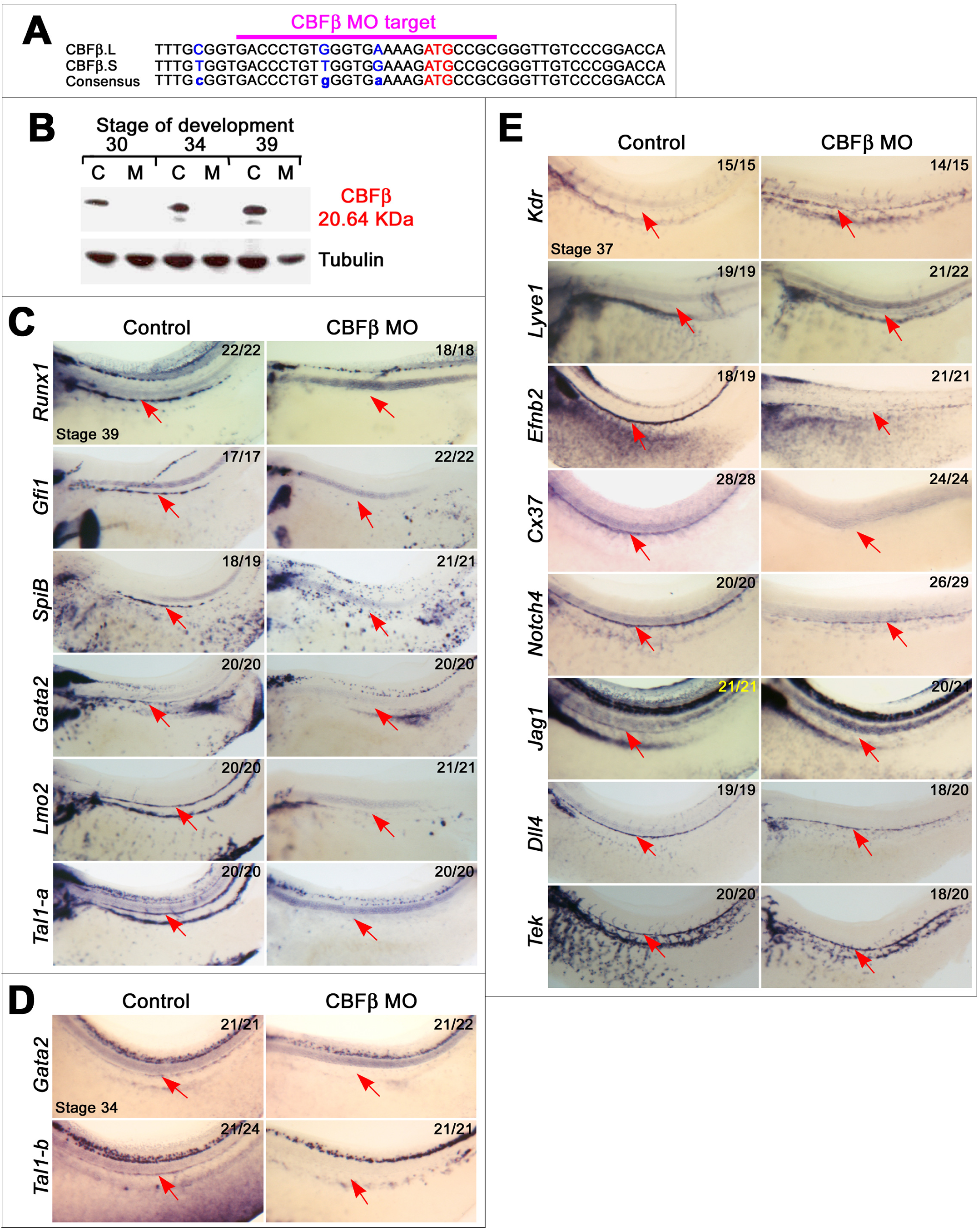
CBFβ regulates HE, arterial and venous expression in the DA. (**A**)Schematic showing CBFβ MO target design. (**B**) Western blot showing that CBFβ protein is absent in CBFβ morphants. Whole embryos were blotted at the stages indicated. C, control; M, MO injected (35ng). Tubulin was used as a loading control. (**C-D**) ISH showing that expression of HE (**C**) and blood genes expressed before the onset of Runx1 (**D**) is absent in the DA of CBFβ morphants. (**E**)ISH showing that, with the exception of Dll4, the expression of the arterial genes (Efbn2, Gja4, Notch4 and Jag1) is downregulated in the DA of CBFβ morphants whereas Kdr expression fails to be repressed. Arrows in (**C**-**E**) indicate expression in the DA. Numbers in top right corner indicate proportion of embryos displaying the phenotype.

### CBFβ is downstream of BMP and regulates VEGFA signaling to program DH

To elucidate CBFβ’s function, we analysed its expression pattern before the formation of the DA and found that it is expressed in the LPM (Figure 5A), where its expression is reminiscent of that of Tal1, a TF which defines the DH identity (Ciau-Uitz et al., 2013). Importantly, expression analysis showed that Tal1 is absent in the LPM of CBFβ morphants (Figure 5B), demonstrating that CBFβ is required for DH specification. To further understand CBFβ’s mechanism of action, the gene regulatory network (GRN) controlling DH establishment was analysed in CBFβ morphants. We have previously demonstrated that VEGFA signaling in the LPM is required for Tal1 expression and that its expression is controlled by Etv6 (Ciau-Uitz et al., 2013). Expression analysis in CBFβ morphants showed that Etv6 expression in the LPM is dependent on CBFβ and, consequently, VegfA is not expressed in these embryos (Figure 5B). In addition to the LPM, both Etv6 and VegfA are expressed in the somites but their expression in this tissue is not affected when CBFβ is depleted, which is in agreement with the lack of CBFβ expression in the somites (Figure 5A). Etv6 expression in the LPM is dependent on Kdr activation by its ligand, VEGFA, emanating from the somites and, therefore, its expression is absent in Kdr deficient embryos (Ciau-Uitz et al., 2010). CBFβ expression is also dependent on Kdr and is not expressed in the LPM when VEGFA is depleted (Figure 5C), this indicates that CBFβ expression in the LPM is dependent on VEGFA signaling from the somites and that it is hierarchically upstream of Etv6.

**Figure 5.**
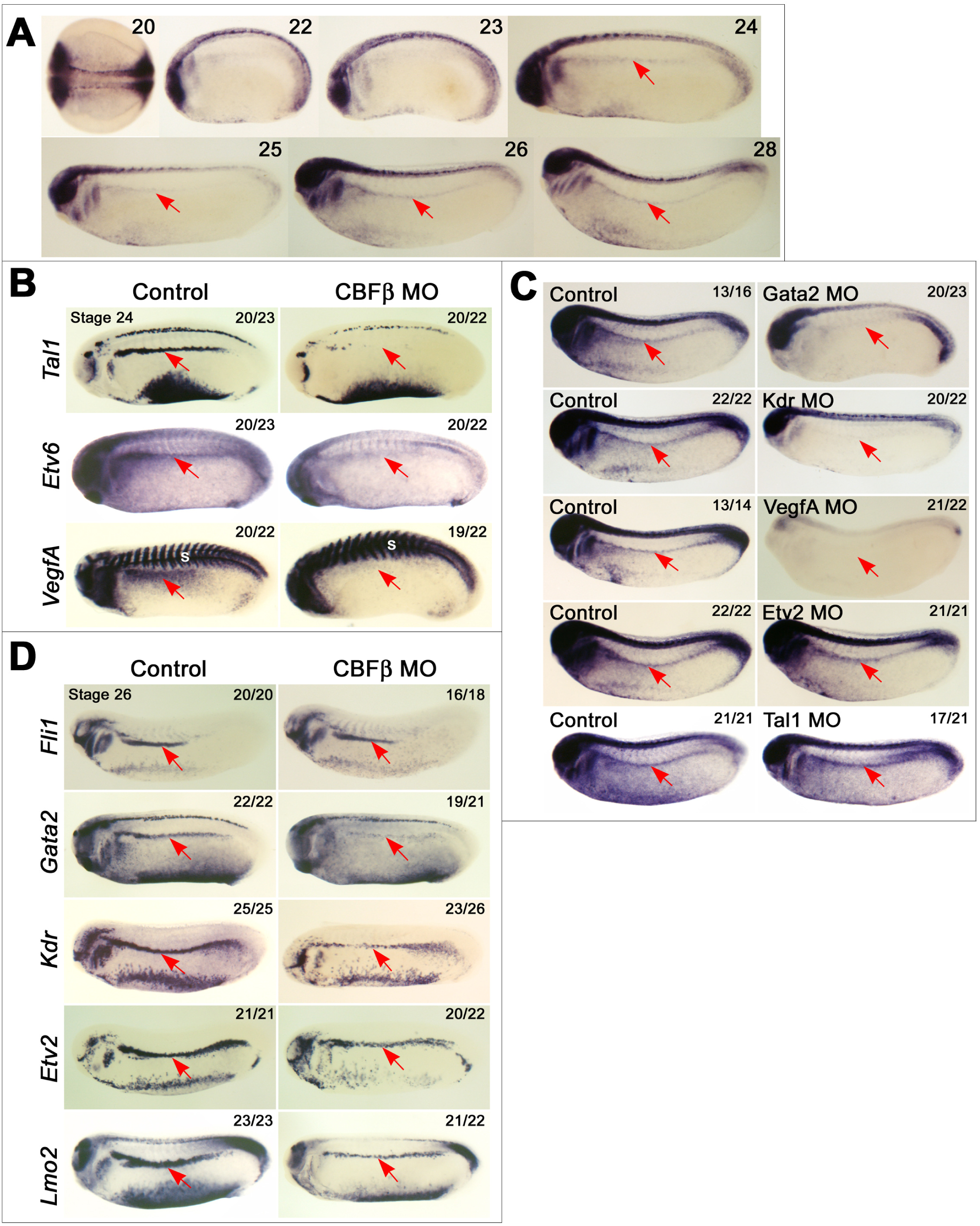
CBFβ is required for DH specification in the LPM. (**A**)ISH showing expression of CBFβ in the LPM (arrow). Numbers in top right corner indicate stage of development. (**B**) ISH at stage 24 showing that CBFβ is required for Tal1, Etv6 and VegfA expression in the LPM. (**C**)ISH at stage 26 showing that CBFβ expression is absent in the LPM of Gata2, Kdr, VegfA and Gata2 morphants but unaffected in Etv2 and Tal1 morphants. MO concentrations as previously indicated (Ciau-Uitz et al., 2013). (**D**)ISH at stage 26 showing that Gata2, Kdr, Etv2 and Lmo2 expression is downregulated in the LPM of CBFβ morphants whereas Fli1 expression is unaffected. Arrows in (**B-D**) indicate expression in the LPM. Numbers in top right corner indicate proportion of embryos displaying the phenotype.

Although CBFβ expression in the LPM is dependent on Kdr, Kdr itself is downregulated in stage 26 CBFβ morphants (Figure 5D), suggesting that CBFβ plays an earlier role that is independent of VEGFA. We have previously showed that LPM expression of Fli1, Gata2, Etv2 and Lmo2 is independent of VEGFA (Ciau-Uitz et al., 2013); strikingly, the expression of these genes, but not Fli1, was also downregulated in CBFβ morphants (Figure 5D). DH programing begins when Fli1 is activated in the LPM, then Fli1 activates Gata2 and together activate Kdr and Etv2 (which maintains Gata2 and activates Lmo2)(Ciau-Uitz et al., 2013). However, Fli1 alone can partially activate Kdr and Etv2 at levels which are insufficient for Tal1 expression. Nevertheless, their levels of expression is enough for the survival of Kdr cells which later migrate to form the DA but do not become HE because they were not specified as DHs. This suggests that CBFβ acts upstream of Gata2 but downstream of Fli1.

To confirm CBFβ’s position in the DH GRN, we analysed the expression of Fli1, Gata2, Etv2 and Kdr at stage 22, when the GRN starts to be established. At this stage, Gata2, Etv2 and Kdr are already downregulated in the LPM of CBFβ morphants whereas Fli1 appears unaffected (Figure 6A), thus, confirming that CBFβ regulates DH specification through Gata2. The CBFβ phenotype is reminiscent of BMP signaling inhibition (Kirmizitas et al., 2017). Furthermore, CBFβ has recently been identified as a target of BMP4 signaling in breast cancer cells (Ampuja et al., 2016) and is required for the proper activation of BMP signaling in chondrocytes (Park et al., 2016). Therefore, we investigated whether BMP controls CBFβ expression in the LPM. To this aim, embryos carrying a heat shock inducible Noggin transgene (Beck et al., 2006) were heat shocked at stage 19 and gene expression was analysed at stage 26. Inhibition of BMP signaling by Noggin resulted in the abrogation of CBFβ, Gata2, Etv6, VegfA and Tal1 expression in the LPM (Figure 6B), indicating that BMP signaling regulates DH specification through CBFβ. Remarkably, as in CBFβ morphants, Etv6 and VegfA expression in the somites was unaffected in BMP deficient embryos, indicating that a BMP/CBFβ independent mechanism controls their expression in this tissue. Also, these results confirm our previous finding that VEGFA signaling from the somites is not sufficient for DH specification but that the establishment of endogenous VEGFA signaling is required (Ciau-Uitz et al., 2010).

**Figure 6.**
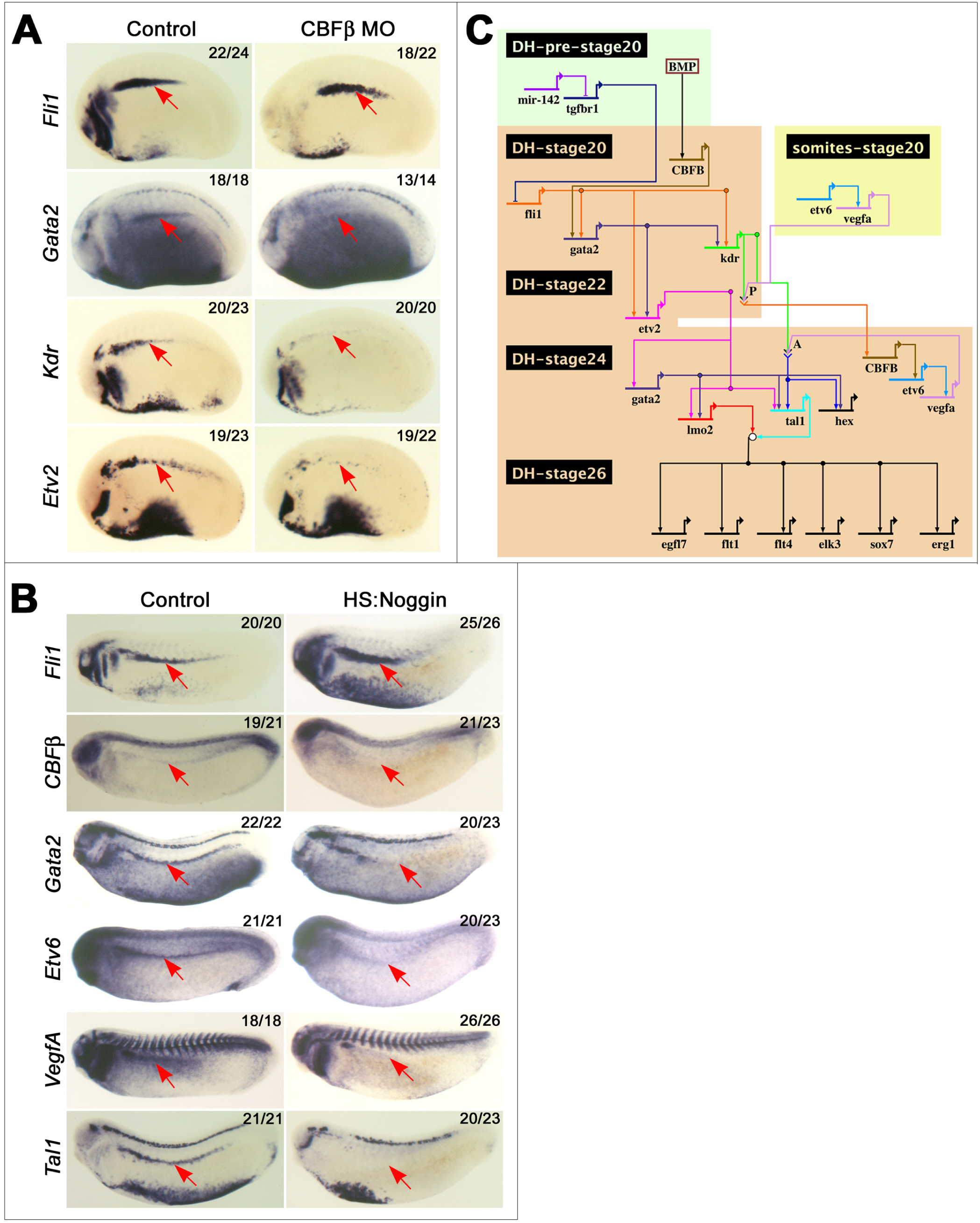
CBFβ mediates BMP signaling to specify DH in the LPM. (**A**) ISH at stage 22 showing that expression of Gata2, Kdr and Etv2 but not Fli1 is downregulated in the LPM (arrows) of CBFβ morphants. Numbers in top right corner indicate proportion of embryos displaying the phenotype. (**B**)ISH at stage 26 showing the absence of CBFβ, Gata2, Etv6, VegfA and Tal1 expression in the LPM (arrows) of HS:Noggin embryos. Embryos were heat shocked at stage 19 as previously described (Kirmizitas et al., 2017). Numbers in top right corner indicate proportion of embryos displaying the phenotype. (**C**)Updated GRN showing the inputs of CBFβ in the programing of DH in the LPM.

Our current knowledge of the GRN controlling DH specification (Ciau-Uitz et al., 2013; Kirmizitas et al., 2017; Nimmo et al., 2013), including the inputs of CBFβ, is presented in Figure 6C. In brief, initially, CBFβ mediates BMP signaling to activate Kdr which makes the LPM precursors responsive to VEGFA ligand emanating from the somites. Then, CBFβ becomes dependent on this paracrine VEGFA signalling and, critically, establishes endogenous VEGFA signaling through the activation of Etv6. This leads ultimately to the activation of Tal1 expression and the emergence of DH.

### CBFβ programmes DH independently of its interaction with Runx

CBFβ function in the establishment of definitive hematopoiesis is thought to be dependent on its binding to Runx1. Here, we have shown that CBFβ is essential for the specification of DH in the LPM, a tissue where Runx1 is not expressed (Figure 1A)(Ciau-Uitz et al., 2013). In agreement, DH specification in Runx1 deficient embryos is unaffected (Figures 7A and S2F). As CBFβ transcriptional activity is thought to be strictly dependent on its binding to Runx proteins, we investigated whether Runx2 or Runx3 are expressed in the LPM. It has previously been reported that Runx2 is undetectable by RT-PCR in whole embryos at stage 25 (Kerney et al., 2007). In agreement, ISH analysis showed that Runx2 is not expressed at the time DH is specified in the LPM (Figure 7B). Similarly, Runx3 expression has not been reported in the Xenopus LPM (Park and Saint-Jeannet, 2010) and our ISH analysis shows that it is exclusively expressed in the somites (Figure 7C). Therefore, we conclude that no Runx gene is expressed in the LPM at the time CBFβ specifies DH.

**Figure 7.**
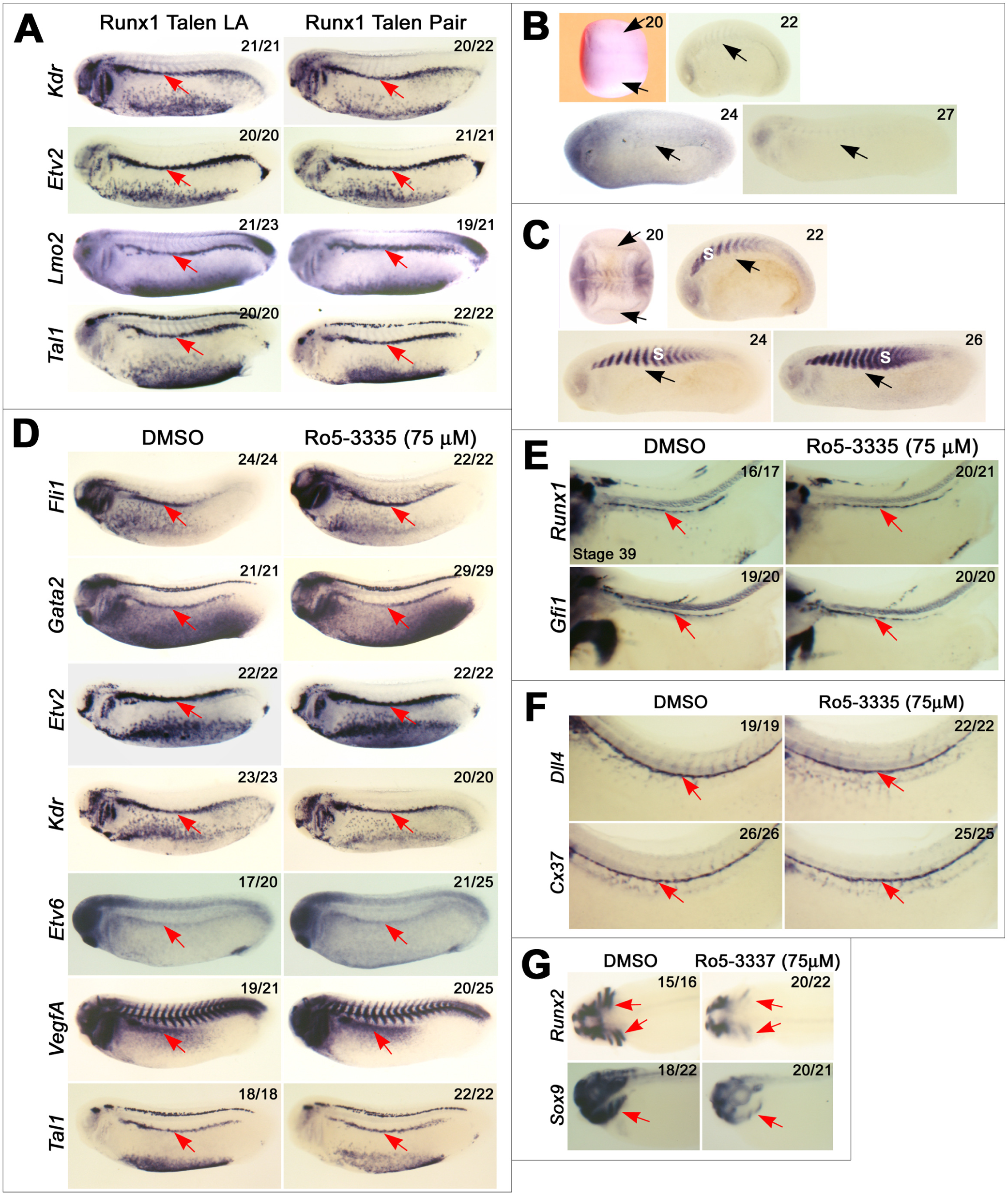
CBFβ regulates gene expression in the LPM independently of Runx proteins. (**A**)ISH at stage 26 showing that DH specification in the LPM (arrow) is unaffected in Runx1 deficient embryos generated by TALEN injection. (**B**)ISH showing that Runx2 is not expressed in the LPM (arrows). 20, dorsal view; 22-27, lateral view. Anterior to the left. Numbers in top right corner indicate stage of development. (**C**)ISH showing that Runx3 is not expressed in the LPM (arrows) but in the somites (s). Orientation and numbers as in **B**. (**D**) ISH at stage 26 showing that treatment with Ro5-3335, a drug which blocks CBFβ binding to Runx proteins, does not affect DH specification in the LPM (arrows). (**E**) ISH at stage 39 showing that treatment with Ro5-3335 does not affect HE specification in the DA (arrows). (**F**) ISH at stage 37 showing that treatment with Ro-3335 does not affect arterial expression in the DA (arrows). (**G**) ISH at stage 39 showing that head cartilage (arrows) development is impaired in Ro5-3335 treated embryos. Numbers in top right corner (**A**, **D**-**G**) indicate proportion of embryos displaying the phenotype.

To test whether CBFβ gene regulation activity can be independent of its binding to Runx, we decided to block their physical interaction using a small molecule. Several small molecules have been designed to disrupt CBFβ binding to Runx (Cunningham et al., 2012; Illendula et al., 2016) and one, Ro5-3335, has been successfully used in developing zebrafish embryos (Bresciani et al., 2014). Therefore, as DH programming initiates at stage 20, we treated embryos with Ro5-3335 from stage 17. If CBFβ‘s transcriptional activity is dependent on its binding to a Runx protein, this treatment would phenocopy CBFβ deficiency. However, expression analysis shows that treatment with Ro5-3335 has no effect on DH specification (Figure 7D), indicating that CBFβ regulates gene expression in the LPM independently of Runx proteins. Additionally, Ro5-3335 treatment did not affect HE establishment in the DA (Figure 7E) because the Runx1-CBFβ interaction is not required for this process. Similarly, expression of arterial genes in the DA is not altered by Ro5-3335 (Figure 7F). It has been shown that CBFβ plays an essential role in chondrocyte differentiation by binding to, and stabilising, Runx2 and Runx3 (Park et al., 2016). As expected, head cartilage development was impaired in Ro5-3335 treated embryos (Figure 7G), indicating that this drug is indeed actively blocking the physical interaction between CBFβ and Runx proteins in Xenopus embryos.

In conclusion, we have discovered that CBFβ plays an essential role in the initiation of the HSC lineage and that this role is independent of its binding to Runx proteins. In other words, we have uncovered that CBFβ’s transcriptional activity is not strictly dependent on Runx.

## DISCUSSION

Runx1 marks HE in the ventral wall of the DA and plays an essential role in the generation of HSCs by regulating EHT. To better understand how HE is established, we profiled gene expression in the DA before and after the onset of Runx1 expression up to the budding of the first IACs (Figure 1C). This led to the following observations: (a) initially, the DA expresses both arterial and vein affiliated genes but the vein program, including Kdr expression, is repressed as Runx1 is activated. (b) Runx1 expression is activated concomitantly with DA lumenization but before blood circulation initiates. (c) Gata2 and Tal1-b are expressed before Runx1, suggesting that they regulate Runx1 as reported in other systems. (d) CBFβ expression is activated concomitantly with the formation of IAC. (e) Runx1 direct transcriptional targets are expressed before CBFβ expression. How these processes influence each other is poorly understood. Nevertheless, here we have shown that, at the early stages of HE establishment, Runx1 exclusively regulates hematopoietic expression without affecting arterial or vein programming. It is also important to note that Runx1 expression initiates before the onset of blood circulation, indicating that shear stress does not induce the emergence of HE.

Studies in differentiating ES cells have suggested that Runx1 represses Kdr expression in HE (Hirai et al., 2005). However, here we have demonstrated that Kdr repression in the DA is Runx1 independent (Figures 2D & S3F). This indicates that gene regulation in ES cell-derived HE is different to that of the DA. Another discrepancy between ES-cell derived hematopoiesis and definitive hematopoiesis in the embryo concerns their hemangioblast precursors. It has been reported that in ES cells Runx1 is expressed by, and required for, the specification of hemangioblasts (Lacaud et al., 2002). In contrast, we have shown that Runx1 is not expressed nor required for the specification of DH in the LPM (Figures 1A, 7A & S2F), the precursors of the DA and HSCs in the embryo. Runx1 is, however, expressed in hemangioblasts giving rise to embryonic (yolk sac-type) blood (Walmsley et al., 2002).

Furthermore, while HE in the DA emerges from arterial endothelium, it has been suggested that definitive HE is not related to the arterial lineage in cultured human pluripotent stem cells (Ditadi et al., 2015). Importantly, it has been reported that yolk sac HE can derive from venous rather than arterial endothelium (Frame et al., 2016). Taken together, these strongly suggest that hematopoiesis in ES and iPS systems do not represent DA but rather yolk sac-type hematopoiesis (Ivanovs et al., 2017). Therefore, conclusions regarding HSC development derived from these systems should be drawn with caution.

Loss of function analysis in the mouse has demonstrated that Runx1 and CBFβ are required for the emergence of HSCs. Because CBFβ is a non-DNA-binding protein whose functions are thought to depend entirely on its physical interaction with Runx, it has been believed that the two TFs regulate the same processes during HSC emergence. However, our study demonstrates that these proteins have independent functions and that they regulate distinct processes in HSC ontogeny. First of all, we have demonstrated that CBFβ is not ubiquitously expressed and that it is absent from the DA when Runx1 establishes the HE program. This may indicate that Runx1 interaction with other TFs expressed in the DA, for example Ets1 (Figure 1C)(Shrivastava et al., 2014), is sufficient for its stabilisation and transcriptional activities. Previous reports support the notion that Runx proteins may not always require CBFβ to regulate gene expression. In breast cancer cells, CBFβ is required for the expression of only a subset of Runx2 target genes (Mendoza-Villanueva et al., 2010), indicating that Runx2 can activate gene expression without it. The presence of HSPCs in zebrafish embryos treated with Ro5-3335 has suggested that Runx1 may regulate HSC emergence without CBFβ (Bresciani et al., 2014). Through genetic evidence, we have demonstrated that Runx1 alone is indeed sufficient for the establishment of HE.

Critically, we have demonstrated that CBFβ regulates the establishment of DH independently of Runx proteins. We provide, for the first time, compelling evidence indicating that CBFβ function in vertebrates, in general, and in developmental hematopoiesis, in particular, is not strictly dependent on its physical interaction with CBFα. Because CBFβ functions are thought to be mediated entirely by CBFα, studies on its mechanism of action have been neglected. Nevertheless, studies in C. elegans have suggested that CBFβ may play roles which are independent of Runx (Kagoshima et al., 2007). In this model, the overexpression of either the CBFβ homologue, BRO-1, or the only Runx gene in C elegans, RNT-1, induce the overproduction of seam cells; however, the overexpression of BRO-1 in a RNT-1 null background can still induce seam cell hyperplasia (Kagoshima et al., 2007). This indicates that BRO-1 induced seam cell overproduction is not dependent on RNT-1. Taken together, we believe that there are sufficient grounds to argue that CBFβ should be studied in its own right.

## EXPERIMENTAL PROCEDURES

### Morpholinos and TALENs

All MOs were obtained from GeneTools LLC (Corvallis, OR, USA). The Runx1 gene generates two main transcripts through the usage of two distinct promoters (P1 and P2), specific MOs targeting the products of both promoters were designed (Figure S2) and injected in combination (40ng/embryo in total, 20ng/embryo each). A MO targeting the translation initiation of CBFβ was designed and injected at a concentration of 35ng/embryo (Figure 4). All other MOs are as indicated previously (Ciau-Uitz et al., 2013). A two-step Golden Gate assembly method using the Golden Gate TALEN and TAL effector kit 2.0 (Addgene) was used to construct the TALEN plasmids containing the homodimer-type FokI nuclease domain (Cermak et al., 2011). 2 TALEN pairs targeting the Runx1 locus were designed using the online design tool, Mojo Hand (http://www.talendesian.org/). TALEN mRNA for injection was generated by linearizing with Not1 and transcribing with SP6 RNA polymerase using the mMESSAGE mMACHINE Kit (Ambion). TALEN mRNAs were injected at the 2-cell stage or in the ventral marginal zone (VMZ) of 4-cell stage embryos. For further details of TALEN design, targeting sites, validation and genotyping see Figure S3.

### Embryo manipulation, in situ hybridization (ISH) procedures and Western blot analysis

Xenopus laevis embryos were obtained and cultured as described (Walmsley et al., 2005) and staged according to (Nieuwkoop and Faber, 1967). A heat shock-inducible Noggin transgenic line (Beck et al., 2006) was used to generate BMP-deficient embryos; embryos were heat shocked and processed as previously indicated (Kirmizitas et al., 2017). Ro-3335 (Calbiochem) was dissolved in DMSO and treatments were performed at 75 μM, with DMSO as a vehicle. Embryo microinjection and whole mount ISH was performed as previously described (Walmsley et al., 2005). Before photography, embryos were cleared in benzyl benzoate:benzyl alcohol (2:1). ISH to sectioned embryos was as previously indicated (Ciau-Uitz et al., 2000). For probe details see Table S1. Probes have been deposited into the European Xenopus Stock Centre, University of Portsmouth, UK. Protein extraction and western blot analysis were as previously described (Afouda et al., 2005); CBFβ protein was detected using an anti-Human CBFβ antibody (abcam ab33516), at a 1:1000 dilution.

### Cloning and rescue experiments

cDNAs for wild-type mRunx1 as well as the mRunx1 mutants, R174Q and T161A, were kindly provided by Nancy Speck (Roudaia et al., 2009). To generate constructs for tol2 mediated transgenesis, mRunx1 cDNAs were HA-tagged and cloned into the mini-tol2 vector under the control of the EF1α promoter (Urasaki et al., 2006) using the primers indicate in Table S2. mRunx1 Y113A/T161A double mutant was generated by performing PCR-mediated mutation on the T161A mutant using the primers indicated in Table S2. For rescue experiments, mini-tol2 vectors, Tol2 transposase mRNA and Runx1 MOs we co-injected into the VMZ of 4-cell stage embryos and then cultured to the desired stages of development. Figures were prepared using Photoshop. All animal work was carried out according to UK Home Office regulations under the appropriate project license.

## Footnotes

**AUTHOR CONTRIBUTIONS** Conceptualization, A.C-U. and R.P.; Methodology, A. C-U., P.P., C.F. and A.K.; Investigation, A.C-U., P.P., C.F. and A.K.,; Visualization, A.C-U., Supervision, A.C-U. and R.P.; Writing-Original draft, A.C-U.; Writing-Review & Editing, P.P., A.K., A.C-U., and R.P. Funding Acquisition, R.P.

## ACKNOWLEDGEMENTS

We are grateful to Nancy Speck for providing the mouse Runx1 DNAs (WT and R174Q and T161A mutants). This work was supported by the UK MRC. We thank Rossella Rispoli for drawing the DH GRN.

Authors declare no conflict of interest.

